# Metabolic constraints on growth explain how developmental temperature scales synaptic connectivity relevant for behavior

**DOI:** 10.1101/2023.10.15.562389

**Authors:** P. Züfle, L. L. Batista, S.C. Brandão, G. D’Uva, C. Daniel, C. Martelli

## Abstract

Environmental temperature dictates the developmental pace of poikilothermic animals. In *Drosophila*, brain development at lower temperature is not only slower, but it also results in different wiring outcomes. A first-principle model that imposes different metabolic constraints for the growth of the neural system and the organism explains these findings, predicts brain wiring under ecologically relevant temperature cycles and explains the non-uniform scaling of neural development across temperatures. Dissecting the circuit architecture and function of first, second and third order neurons in the olfactory system, we demonstrate that the consequences of temperature are contingent upon the availability of synaptic partners in different circuits. Despite synaptic scaling, second order neurons encode robust odor representations, while temperature dependent connectivity of third order neurons leads to differences in odor-driven behavior. Therefore some circuit specific developmental programs have evolved to support functional robustness with respect to environmental temperature, while others allow phenotypic plasticity with possible adaptive advantages.

## Main Text

The wiring of the nervous system follows a complex genetic plan during development. However, due to stochastic processes and environmental factors, genetically identical individuals seldomly show the same phenotypic outcome^1^. Temperature is the environmental factor with the broadest effects in biology, as it determines the rates of all biophysical reactions of an organism ^2^. In poikilothermic animals, such as insects, worms, fish, amphibians, and reptiles, temperature determines the speed of development. Mathematical theories of growth have shown that developmental times scale exponentially with temperature due to a constraint imposed by the rate limiting metabolic reaction^3^. The development of the nervous system is certainly not exempted by the effects of temperature.

It has been widely reported that development at different temperatures correlates with variation in behavioral phenotypes with examples in amphibians^4^, reptiles^5–7^, bees^8,9^, ants^10^, and fruit flies ^11,12^. Social insects invest a lot of resources to keep their broods at the correct temperature throughout daily and seasonal cycles. Even small variations in developmental temperature can affect learning in bees ^13^ and the synaptic organization of key brain areas for learning^12,14,15^. Therefore, temperature does not just determine the speed, but also the outcome of development, although the mechanistic bases of this phenotypic variation are largely unknown.

A recent study in *Drosophila* reported that the number of synaptic connections between neurons of the visual system inversely correlate to temperature, i.e. when flies develop at lower temperature these neurons make more synapses and have more postsynaptic partners ^16^. This leads to the question whether synaptic scaling occurs similarly throughout the brain and what consequences it has on neural computations and behavior. Answering these questions is key not only to predict the consequences of temperature changes on animal behavior in the wild, but also to fully understand the organization of development in poikilothermic animals.

### Developmental temperature differentially affects the wiring of individual olfactory glomeruli

To investigate the effects of temperature on the development of similar, but functionally distinct neural circuits, we focus on the olfactory system of *Drosophila*. Olfaction mediates key behaviors in animals, including foraging and mating, plays a major role in the organization of animal societies and mediates adaptation to the environment. We set out to investigate the effect of developmental temperature on the wiring of Olfactory Receptor Neurons (ORNs) within the Antennal Lobe (AL), the main olfactory area in the insect brain. We used trans-Tango^17^, a genetic tool for transsynaptic labeling, to analyze post-synaptic neurons of or42b-ORNs, i.e. ORNs that express the odorant receptor or42b and target the glomerulus DM1 (Fig.1a). Flies developed at either 18°C or 25°C between the larval stage L3 and the end of pupal development (which we name P-100% as the absolute developmental time depends on temperature). Flies were then kept at 25°C for 10 days before dissection. Strikingly more postsynaptic neurons are labeled in flies that developed at 18°C, as suggested by denser innervations in both the AL and downstream areas (Fig.1b-c). We quantified these differences by counting the cell bodies of the postsynaptic cells in each fly, which were more than double at 18°C as compared to 25°C (Fig. 1d).

**Figure 1.**
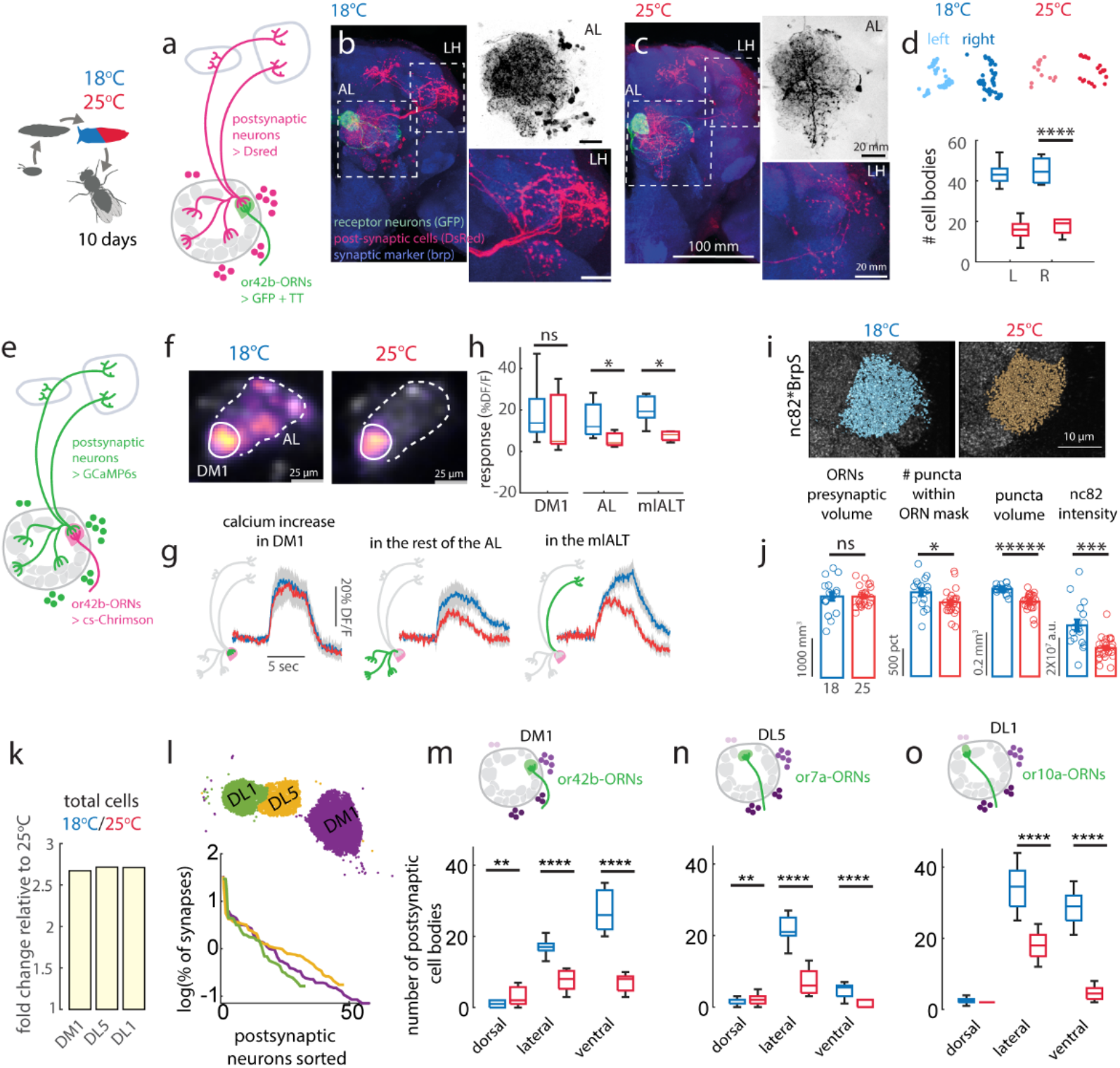
Developmental temperature affects the connectivity of ORNs in the antennal lobe. **a)** Flies expressing the trans-Tango construct under control of or42b-Gal4 were placed at 25°C or 18°C between L2 and P-100% and then dissected when 10 days old. Or42b-ORNs send its axons in the DM1 glomerulus where they target both local neurons (LNs) and projection neurons (PNs) that send axons to higher brain regions. **b-c)** Example brains from flies developed at the two temperatures. Immunostaining labels ORNs that express GFP (green), post-synaptic partners expressing dsRed (red) and all synapses through anti-Brp (nc82, blue). Insets are close-ups of the innervations in the AL (dsRed signal in grey scale) and LH. **d)** Cell body locations in left (L) and right (R) hemisphere in two individual flies and boxplot showing median and quartiles of the number of postsynaptic partners, Kruskal-Wallis test, p < 10^−7^, n=10. **e)** Schematics of the experiment where the usual trans-Tango reporter dsRed was replaced by GCaMP6s, and activity of presynaptic neurons is induced by CsChrimson. **f)** Example responses for flies developed at different temperatures. Color bar indicates ΔF/F calculated for each pixel. White lines indicate the region of interests (roi) used for quantification. **g)** Optogenetic response to 0.05mW/mm^2^ quantified in the DM1 glomerulus, in the rest of the AL and in the mlALT. **h)** Boxplot showing median, quartiles and min/max response taken as the mean in 9s following stimulus onset. Significant differences were observed in the AL roi (Kruskal-Wallis test, p=0.02, n=4-7) and in the mlALT (Kruska-Wallis test, p=0.01, n=4-7). **i)** nc82 staining of synaptic puncta and 3D reconstruction of the puncta within the ORN mask created from Brp^[Short]^-GFP fluorescence for the two temperatures. **j)** Volume of the Brp^[Short]^-mask, number of puncta, puncta volume and intensity within the ORN mask (error bars indicate SEM, n = 16 hemibrains at 18 °C and n = 25 at 25 °C, Kruskal-Wallis test, *p<0.05, **p<10^−2^, ***p<10^−3^, *****p<10^−5^). **k)** Fold change in number of synaptic partners relative to 25°C for three glomeruli. **l)** Allocation of ORN output synapses onto individual partners for the three glomeruli, as quantified from EM data (Hemibrain). The number of synapses was normalized to the total for each ORN class. Postsynaptic neurons are sorted based on the number of synaptic inputs they receive. Top: spatial location of the synapses analyzed (only right hemisphere). **m-o)** Trans-Tango analysis of connectivity of three different ORNs. Number of synaptic partners in different cell clusters of the AL. Boxplots indicate median and quartiles, whiskers indicate maximum and minimum values. Sample size at 18/25 °C: DM1 n=20/23, DL5 n=10/10, DL1 n=10/10, Kruskal-Wallis test *p<0.05, **p<10^−2^, ***p<10^−3^, *****p<10^−5^.

To demonstrate that the anatomically labelled postsynaptic neurons are functionally connected to the ORNs, we modified the trans-Tango experiment to express a calcium reporter (GCaMP6s) in the post-synaptic neurons, and CsChrimson in or42b-ORNs (Fig.1e). This allowed us to activate optogenetically specific ORNs and test response in their postsynaptic partners. Electrophysiological experiments indicate that optogenetic activation of ORNs drives less spiking activity than odors (not shown), but sufficient to induce postsynaptic responses in a dose-dependent manner (Supp. Fig. 1c). Calcium transients measured within the DM1 glomerulus mostly report activity from the single uniglomerular PN (uPN, which receives most of the ORN synapses) and do not differ at the two temperatures (Fig. 1g, Supp. Fig. 1a). However, activation in the rest of the AL is higher in flies developed at 18°C (Fig. 1f-h), which is consistent with a larger number of multiglomerular neurons being connected to DM1. Moreover, activity is stronger in the medio-lateral AL track (mlALT, Fig. 1g,h) through which a class of multi-glomerular inhibitory PNs (vPNs) project their axons to the Lateral Horn (LH). Therefore, trans-Tango labeled neurons are functionally connected to the or42b-ORNs, and ORNs drive more activity in flies developed at 18 °C.

More connected neurons could result from a different distribution of synapses across postsynaptic partners or from an increase in synapse number. To distinguish between these two cases, we expressed a GFP tagged Brp^Short^ in or42b-ORNs to label presynapses and stained the endogenous Brp to count synaptic puncta^18^ (nc82, Fig. 1i). The or42b-ORNs presynaptic volume defined by the Brp^Short^ mask was consistent across the two developmental temperatures (Fig. 1j). However, flies developed at 18°C had higher Brp^Short^ intensity, an increased number of nc82-labeled Brp puncta within the presynaptic volume, larger puncta volumes and stronger intensity (Fig. 1j). Both fluorescence intensity and the number of synapses within the whole volume of the DM1 glomerulus, which includes connections between other neuron types, were also higher (Supp. Fig. 1c). Together, these results demonstrate that a lower developmental temperature leads both to more synapses, larger active zones, and more synaptic partners which are functionally connected to the ORNs.

A similar fold change in post-synaptic partners was found across three glomeruli (Fig. 1k). This is consistent with a similar distribution of synaptic connections onto other neurons in these glomeruli, which we quantified from EM data (Fig. 1l, and Supp. Fig. 1d), such that if all connections are scaled up or down by temperature, the relative changes would be similar across glomeruli, as observed. We postulate that despite a similar scaling, the identity of synaptic partners recruited by these different ORNs will depend on available neurons in the relevant glomerulus volume. AL neurons postsynaptic to ORNs have their cell bodies distributed in four clusters^19^. Or42b-ORNs increased connections more with neurons of the ventral and ventrolateral clusters (Fig. 1m) that host cell bodies of 66.6% of all multiglomerular PNs (mPNs)^20^ including vPNs^21–24^. This suggests that Or42b-ORNs more strongly drive this population of neurons when flies develop at 18°C and is consistent with imaging in the mlALT (Fig. 1h). On the contrary, within the glomeruli DL5 and DL1 the corresponding ORNs increase connectivity more strongly with neurons of the lateral cluster (Fig. 1n-o) that contains a large population of LNs, and mPNs mostly of unknown function^25^. EM data confirm that more mPNs innervate the DM1 glomerulus than the DL1 or DL5 glomeruli, where we find instead a larger percentage of LNs, which are mostly inhibitory (Supp. Fig. 1e). These findings suggest different functional consequences of developmental temperature in different glomeruli. Specifically, we predict a stronger recruitment of local inhibition in all glomeruli at lower temperatures. For the DM1 glomerulus, we predict it drives stronger activity in inhibitory mPNs that send axons to the LH through the mlALT. Silencing of inhibitory mPNs from the ventral cluster has been shown to drastically reduce approach behavior to appetitive cues^21^ therefore the observe structural changes have potential functional consequences downstream of the AL.

### A first-principle model explains the scaling of brain wiring at different temperatures

Next we asked how synaptic connectivity scales across a wider range of temperatures. *D. melanogaster* lives in different climates, with a thermal range between 11°C and 31°C^26^. We used trans-Tango to characterize connectivity patterns of or42b-ORNs in flies that developed at extreme temperatures (12-31°C). These conditions are highly stressful, and if persistent, female reproductive success would go to zero^27^. However, flies were viable to temperature shifts restricted to pupal development. In flies developed at 12°C, ORNs established a larger number of synaptic partners, including neurons that target regions outside the canonical olfactory pathway (Fig. 2a). Importantly, the anatomy of the ORNs is strongly affected at this temperature, showing axonal extensions protruding radially from the DM1 glomerulus (Fig. 2c vs 2e) as well as mistargeting to a more anterior glomerulus, VA2 (Fig. 2d). Flies developed at 31°C instead show very few neurons connected to or42b-ORNs (Fig. 2b,f).

**Figure 2.**
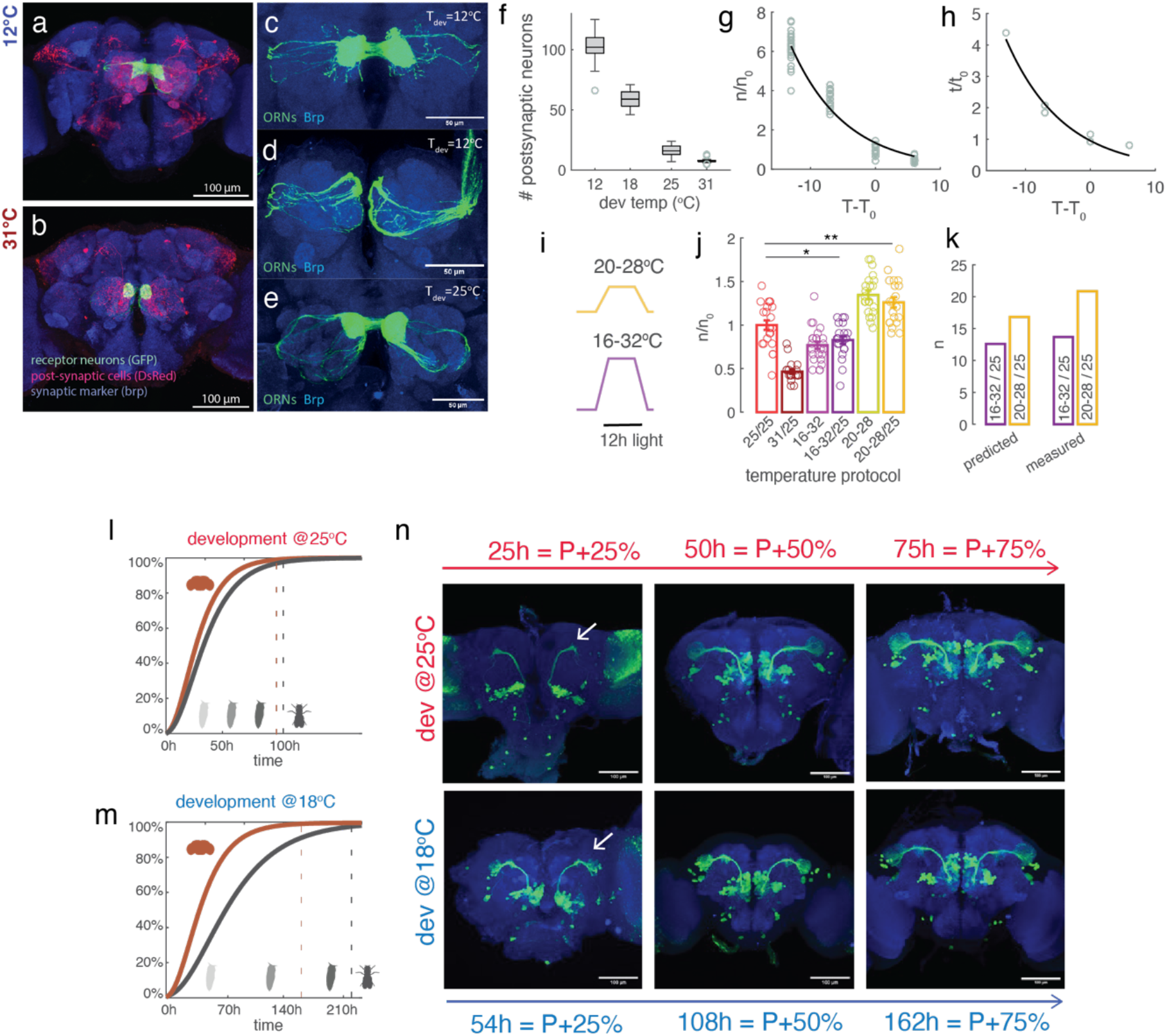
A first principle-model for network connectivity at different temperatures. **a-b)** Example brains of flies expressing trans-Tango under control of or42b-Gal4 and developed at 12°C or 31°C. **c-e)** Magnification of ORN axons labeled by GFP. **c)** Or42b-ORNs mistargeting outside the DM1 glomerulus in flies that developed at 12°C. **d)** Same fly as in **c** imaged at a more anterior focal plane, showing innervations of the or42b-ORNs in the glomerulus VA2. **e)** Normal axonal projections in flies developed at 25°C. **f)** Box plot of the number of postsynaptic partners as a function of developmental temperature. **g)** Single data points as in **f**, normalized by the mean number of neurons connected at a reference temperature (25°C) and plotted as a function of the difference from the reference temperature. The line indicates the exponential fit. **h)** Fold change in developmental time with respect to the temperature difference from 25°C and exponential fit. **i)** Schematics of the temperature and light cycles. **j)** Fold change in synaptic partners from 25°C for all temperature protocols. Temperature shifts were applied only between L2 and P-100% then flies were moved to 25°C, except for the light-purple (16-32) and light-yellow (20-28) bars for which flies were kept through adulthood at the cycling temperatures. **k)** Predicted (see Methods) and measured (same data as in **j**) absolute numbers of post-synaptic partners in temperature cycles. **l-m**) Schematics of the growth model showing % of development as a function of time for the brain and whole animal at two developmental temperatures. Parameters of the growth dynamics are hypothetical. **n**) Staining of PNs labeled by GH146 at three developmental times corresponding to 25%, 50% and 75% of ontogenesis at the two temperatures.

Across all developmental temperatures tested, the number of ORNs remained constant (Supp Fig. 1b), but the number of synaptic partners scaled exponentially with temperature (Fig. 2f-g), similarly to the scaling of developmental time (Fig. 2h). We reasoned that the scaling of these two factors, number of synaptic partners and developmental time, could derive from similar first principles. Gillooly et al. have proposed that the rate of development scales with temperature proportionally to the Boltzmann factor of the rate limiting metabolic reaction^3^. Assuming that flies eclose to a similar mass across temperatures, developmental time (*t*) should scale with temperature (*T*) as 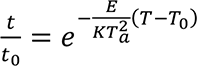 (see Methods), where *t*_0_ is the developmental time at a reference temperature *T*_0_ (here *T*_0_ = 25℃), *E* is highest activation energy in the metabolism (corresponding to the slowest reaction), *K* is the Boltzmann constant, *T_a_* is the water freezing point (273K, see Methods). In our experiments, the scaling factor for the developmental time, defined as 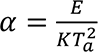, is equal to 0.11 ± 0.02 (Fig. 2h) in agreement with estimates in other animals^3^. This simple exponential model fails to fit developmental times at the highest temperatures above 30°C^28^, nonetheless it provides a good starting point for our theoretical framework. We reasoned that if the development of the neural circuit was limited by the same reaction rate as for ontogenesis (development of the whole organism), then development should result in the same brain connectivity at all temperatures. If, otherwise, the development of the neural system is limited by a reaction with lower activation energy *E*\*, it should scale with the Boltzmann factor 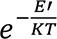 with *E*\* < *E*; the neural circuit will therefore develop (extend axons, form synapses) at its own faster pace, but for an amount of time that is determined by ontogenesis, therefore leading to a larger number of synapses. To calculate what the fold change in connectivity would be, we assumed that axonal growth and synaptogenesis scale with a ¾ power law: 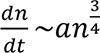 (*n* indicating a measure of neuronal mass and synapses, but see Methods), as for body mass^3^. Many biological processes, including growth, scale allometrically with the ¾ law^29,30^, a phenomenological observation that has received a first principles explanation based on the fractal nature of the distribution of nutrients and resources in a 3D volume by a space-filling network of branching tubes^31,32^. Given the tree-like structure of axons, along which mitochondria and proteins need to be transported, and of other energy suppliers in the brain (such as trachea and glia), the ¾ law is a reasonable assumption for the growth of the neural system. From these assumptions, it is possible to derive an analytical function for the scaling of synaptic connectivity with temperature: 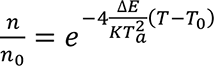 with Δ*E* = *E* − *E*′ (see Methods). We define 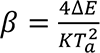 as the scaling factor for synaptic connectivity. When Δ*E* = 0, the same number of connections are established across temperatures. *β* can in principle be different across sub-circuits if their development and growth are differently regulated. In the olfactory circuit we find *β* = 0.12(±0.01). In the visual system, Kiral et al. reported a fold change of ∼1.5 which corresponds to a smaller value of *β*. Whether these differences are technical or biological remains to be determined, but the proposed model would be consistent with differences across circuits.

To test if this first-principle model has any predictive power, we asked what happens in more ecological relevant conditions when flies develop on diurnal temperature cycles. Does development under cycling temperatures result in the same wiring outcome as for flies developed at the mean temperature? We set flies to grow on ecologically realistic temperature cycles with the same mean temperature (24°C) but different amplitudes (20-28°C and 16-32°C, Fig. 2i). These conditions lead to different phenotypes, with more synaptic partners in 20-28°C cycles than 16-32°C cycles, both significantly different from the 25°C condition (Fig. 2j). Further, the temperature experienced after eclosion does not strongly affect connectivity (Fig. 2j). We therefore asked whether the growth model fitted above would be consistent with these findings. Integrating the growth rate over temperature cycles (see Methods) and using the estimate for *α* and *β* from the fixed temperature experiments (Fig. 2g,h), we were able to predict the number of synaptic partners for the cycling temperature protocol (Fig. 2k), and specifically a higher number of synaptic partners for cycles with lower amplitude. The model further predicts that circuit connectivity scales inversely with the amplitude Δ*T* of the temperature fluctuations as: 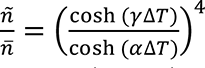, where *ñ* and *n̅* are the number of connections for development on temperature cycles and at the mean temperature, while *γ* depends on both *α* and *β*. Therefore, fluctuating temperatures always lead to less connections than at the mean temperature *T̅*. We conclude that temperature-dependent synaptic scaling is consistent with a growth model with different rate limiting metabolic reactions for the whole organisms and for the brain, with possible differences across neural sub-circuits.

The most general prediction of such model is that while a decrease in temperature leads to both slower ontogenesis and slower brain development, the temporal scaling between these two processes is not the same at the two temperatures (Fig. 2l-m). This implies that the developmental stage of the brain at proportional times of pupal metamorphosis (P-25%, P-50%, P-75%) is expected to be more advanced in flies that develop at 18°C, compared to 25°C. Consistent with this prediction, we find that PNs axons already target the LH at P-25% when flies develop at 18°C, while axonal innervations of the LH are not yet visible at P-25% in flies developed at 25°C (Fig. 2n) consistent with previous data^33^. We conclude that temperature induces a different scaling of brain development compared to ontogenesis, leading to consequences for the wiring of the nervous system.

### Developmental temperature affects odor driven behavior

We next asked whether wiring consequences of developmental temperature affect odor driven behavior. We tested the odor response of individual, 10 days old flies tethered to walk on a spherical treadmill (Fig. 3a). A puff of 2-butanone elicited on average a stronger increase in walking speed in flies that developed at 18°C compared to 25°C (Fig. 3b). These odor responses were highly reproducible across repetitions of the stimulus (Fig. 3d left), although variable across individuals (Fig. 3d right). Similar results were obtained with vinegar (Supp. Fig. 2a-c). Basal walking speed was in general larger for flies developed at 18°C (Fig. 3c). A higher basal walking speed does not justify a higher response, rather the opposite, as flies that walk faster in this assay might not be able to reach a change in speed as high as slower flies.

**Figure 3.**
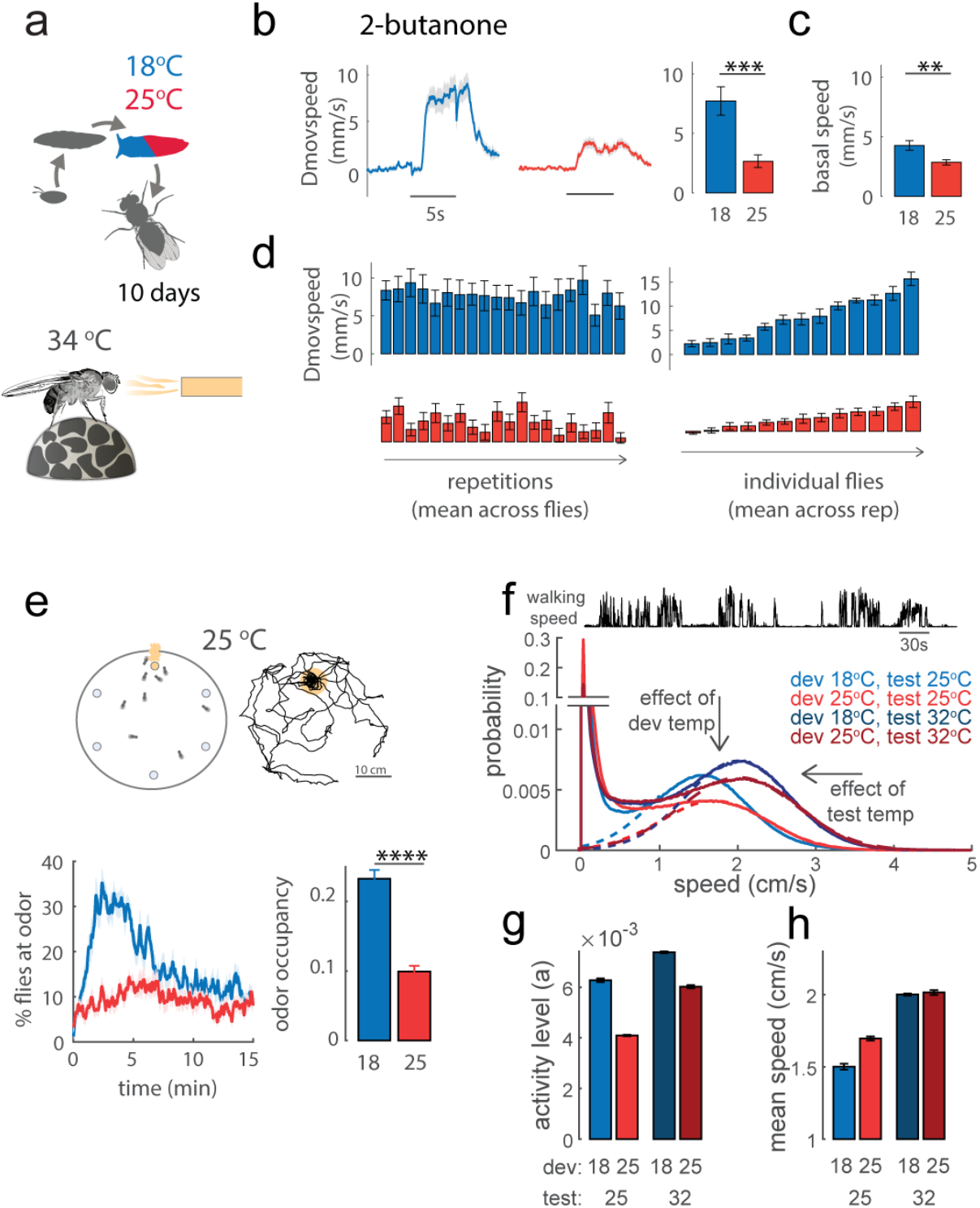
Developmental temperature affects odor-driven behavior. **a)** Flies were placed at 25°C (red) or 18°C (blue) between L2 and P-100% and otherwise kept at 25°C. When 10 days old, odor response was tested at 34°C on a spherical treadmill setup (see Methods). **b)** Odor response calculated as change in moving speed as a function of time in response to 5s stimulation (Δmovspeed = speed - basal speed); bar plot: mean change in moving speed within the first 2s of stimulation (p<10^−3^, n=13/13). **c)** Basal walking speed is estimated within 3s before stimulus onset (p = 0.02). **d)** Mean response to each consecutive stimuli repetitions averaged across all flies (right), and mean response of each individual fly across repetitions, showing large variation across individuals (left). **e)** Top: Same breeding protocol as in **a)**, but odor preference was tested at 25°C in a free walking assay consisting in a circular arena of 40 cm radius with an odor source randomly placed in one of six possible locations. Flies were tracked for 15 min (see Methods). Bottom: Average percentage of flies at the odor source and odor occupancy (bar plot). Shaded area and error bars indicate SEM. Odor occupancy was calculated as the integral under the curve in the first 7 minutes, being 1 if all flies spend 100% of the time at the odor (p<10^−6^, n=19/20). **f)** Top: example of walking speed for one fly, showing bouts of activity. Bottom: probability of walking speed calculated from all data (flies and times) of each condition, showing a bimodal distribution with a large peak in zero that represents inactive flies; dotted lines indicate gaussian fit 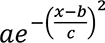 to the active flies. **g-h)** Parameter estimates from the gaussian fit: the amplitude of the gaussian (*a*) as an estimate of the activity levels (probability to be active), and the mean (*b*) as an estimate of the mean walking speed during active times. Error bars indicate confidence intervals on the fitted parameters.

To test whether these findings are specific to this assay, we used a free-walking assay where we tracked the flies’ positions while they explored a large circular arena hiding an odor source. At all times more flies raised at 18°C visited the odor source and spent more time at the odor than flies raised at 25°C (Fig. 3e). Similar results were obtained with vinegar (Supp. Fig. 2d-f) and for a higher testing temperature (Supp. Fig. 2g-h). An analysis of walking speed in control experiments with no odor demonstrates that during active times, the flies’ walking speed is mostly determined by testing temperature (Fig. 3f), while developmental temperature affects the activity level, quantified as the probability to find the flies active (Fig. 3g-h). Since at both testing temperatures the activity levels of flies developed at 18°C are higher, the higher odor occupancy of these flies is not due to inactivity or poor locomotion. We conclude that independently of possible effects on the motor system, flies developed at lower temperatures are more attracted to vinegar and 2-butanone.

### Odor coding in the AL is robust to developmental temperature

We set out to test whether these behavioral phenotypes are due to altered odor representations in the AL and related to temperature-dependent wiring. Extracellular recordings from single sensilla showed that or42b-ORN transduction currents (Supp. Fig. 3a) and firing rates (Fig. 4a) elicited by odor stimuli do not differ in flies developed at different temperatures. Similar results were obtained for or59b-ORNs, innervating glomerulus DM4 (Supp. Fig. 3b-c). Moreover, within the antenna, the number of cell bodies labeled by the or-specific GAL4 line did not differ at the two temperatures (Fig. 4b). We expressed the calcium sensor GCaMP6f in ORNs (labeled by orco-GAL4) and imaged activity in their axon terminals in the AL. The glomeruli DM1 and DM4 showed higher presynaptic calcium responses in flies developed at 18°C (Fig. 4c). Since the number of action potentials and the number of ORNs did not differ with temperature, we conclude that larger calcium transients in these glomeruli reflect the larger number as well as the larger size of synapses being activated (Fig. 1i-j). This raises the possibility that postsynaptic uPNs that send odor information to higher brain areas might be activated more strongly. However, uPNs in all glomeruli tested responded similarly in flies developed at 18°C and 25°C (Fig. 4d). Therefore, odor representations in the output of the AL are robust to changes in developmental temperature.

**Figure 4.**
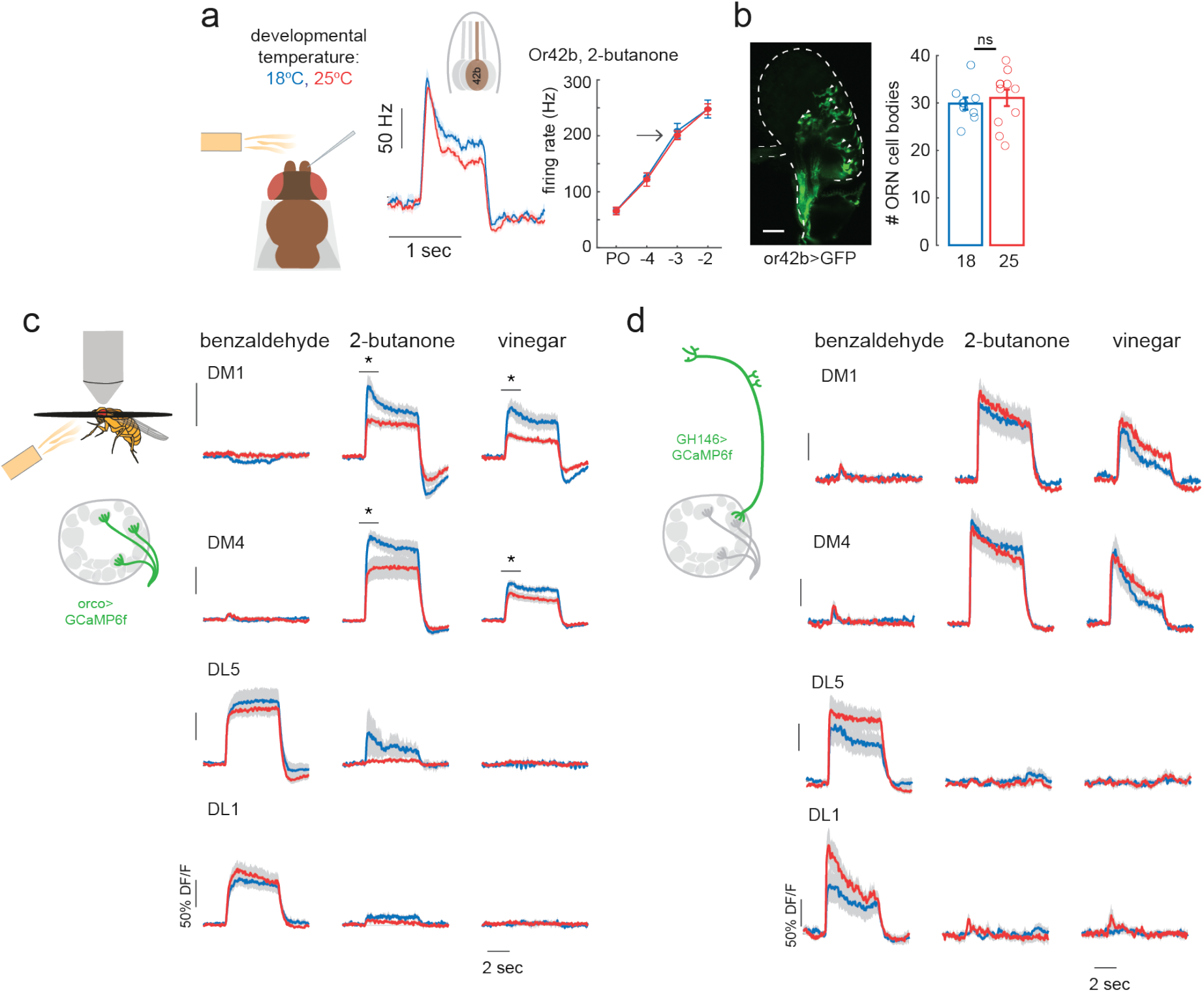
Scaling and normalization of odor responses in input and output neurons of the antennal lobe. **a)** Single sensillum recordings from the ab1 sensillum containing a single or42b-ORN. Left: firing rate response to a 1s pulse of 2-butanone at 10^−3^ dilution calculated in 100ms sliding window. Right: mean peak response for each concentration tested. **b)** Confocal image of the antenna showing GFP expression in or42b-ORNs. Bar plot indicates mean and SEM, individual data points are overlayed. **c)** Calcium imaging from ORN axon terminals in the AL, quantified within three glomeruli for flies developed at the different temperatures (*p<0.05). **d)** Same as **c**, but the calcium reporter was expressed in uPNs and responses were quantified in the dendritic arborizations within corresponding AL glomeruli. Shaded areas indicate SEM, n=5-10 depending on the glomerulus-odor combination.

### Developmental temperature shapes connectivity patterns relevant for innate odor-driven behavior

Since the measured output activity of the AL is invariant to developmental temperature (Fig. 4), we postulated that the different behavioral phenotypes should result from differences downstream of the AL. Innate odor preference is determined by the wiring of uPNs onto LH neurons^34^ and modulated by mPNs^21–24^. We set out to check if the connections of uPNs onto LHNs were also altered by developmental temperature. We labeled the glomeruli connected to a specific LHN type, PD2a1/b1, using the genetically encoded retrograde tracing tool BAcTrace^35^ (Fig. 5a-b). Differently from trans-Tango, this tool only labels presynaptic partners from a pool of genetically labeled neurons, in this case 27 uPNs and 3 mPNs (targeted by VT033006-LexA). Although the innervation of uPNs’ axons into the LH have been shown to be stereotypic across individuals^36^, we found a large degree of inter-individual variation and asymmetry in connectivity across hemispheres (Fig. 5b-c) (also reported by^35^). An analysis of connection probability shows that developmental temperature influences the connectivity pattern of PD2a1/b1 to AL glomeruli, with a stronger effect in the VM2 glomerulus (Fig. 5c-e). Temperature, however, does not affect the number of connected glomeruli (not shown) nor the number of PD2a1/b1 neurons labeled (Fig. 5g). Similarly small, but significant, shifts in connectivity were observed for two other LHNs, AV4a1 and AD1C1 (Fig. 5f, and Supp. Fig. 4).

**Figure 5.**
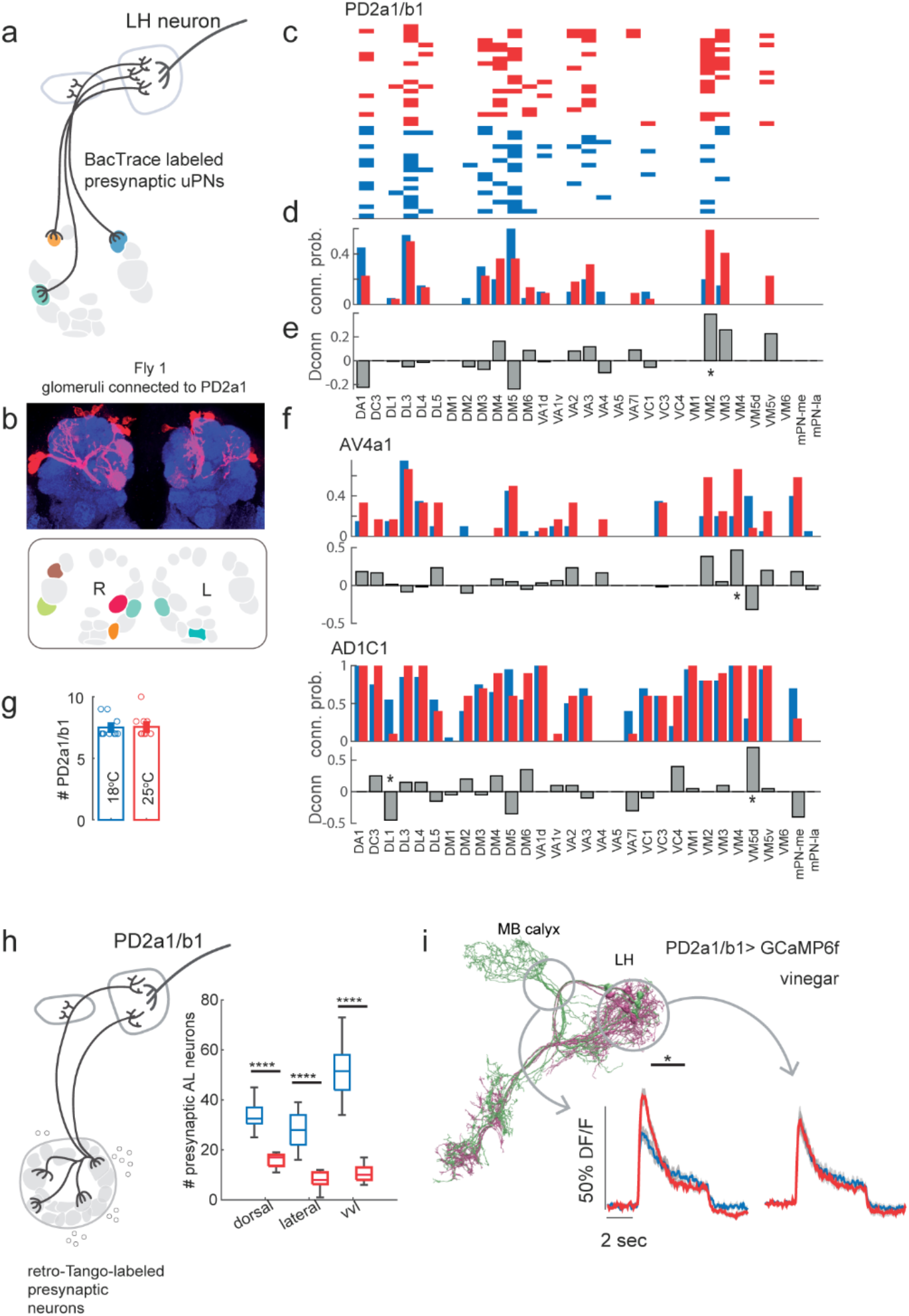
Developmental temperature affects LH neurons connectivity to uPNs and their odor response. **a)** Schematics of the BAcTrace experiment, showing a LH neuron and candidate pre-synaptic uPNs from the AL. **b)** Confocal image of a sample brain showing labeled uPNs arborizations in the AL and binary connectivity patterns of the uPNs (identified by their glomerular innervation) to PD2a1/b1 in a sample brain. **c)** Connectivity matrix between glomeruli (x-axis) and PD2a1/b1 in individual hemibrains (y-axis) in flies developed at 25°C (red) or 18°C (blue). **d)** Probability to find a certain glomerulus connected to PD2a1/b1 for each temperature. **e)** Difference in wiring probability (25°C - 18°C). * = p<0.05, chi-square test, n=22/20. **f)** Difference in wiring probability for two other LH neuron types, AV4a1: n=12/20; AD1C1: n=10/20. **g)** Bar plot showing mean number of PD2a1/b1 neurons labeled by the split-Gal4 line when developed at both temperatures (error bars are SEM, n=12/11 hemibrains). **h)** Schematics of the retro-Tango experiment starting from PD2a1/b1. Boxplot shows median, quartiles and min/max values for the number of PD2a1/b1 presynaptic partners from three AL cell clusters; vvl includes cell bodies in the ventral and ventro-lateral cluster (p<10^−4^, n=12/11 hemibrains). **i**) Calcium responses of PD2a1/b1 to vinegar in flies developed at 25°C (red) or 18°C (blue). Imaging was performed on two focal planes to capture potential differences between the a1 and b1 subtypes, indicated by circles in the EM skeleton. Shaded areas indicate SEM.

The BAcTrace approach restricted the connectivity analysis to uPNs. We therefore used another pan-neuronal retrograde tracer, retro-Tango^37^ to label all the presynaptic partners of PD2a1/b1 (Fig. 5i). Within the AL, we find similar asymmetric labeling of the glomeruli as with BAcTrace (not shown) and a larger number of presynaptic neurons at lower temperatures, with larger effects in the ventral cluster, which contains mPNs (Fig. 5i). This increase in the number of synaptic partners is consistent with a general scaling of connectivity with temperature throughout the brain.

To link these changes in wiring to function, we performed calcium imaging from PD2a1/b1. To separate the a1 and b1 subtypes, we imaged from two focal planes (Fig. 5j). The b1 subtype shows differences in odor response in flies developed at different temperatures (Fig. 5j). This demonstrates that these neurons’ function is not robust to developmental temperature leading to a different LH output in response to odor stimuli. Therefore, while a general scaling of connectivity occurs across the brain, functional consequences are circuit specific and depend on the type of synaptic partners recruited.

## Discussion

The development of a whole organism requires the parallel and coordinated growth of mass and, in the brain, the establishment of functional synaptic connections in different subcircuits. Developmental programs are highly temporally structured, raising the question of how they are affected by a temperature dependent change in developmental speed. Here we propose that developmental speed is set by different biophysical processes in different cell types, with disrupting effects for the temporal coordination of development at different temperatures. This temporal mismatch affects overall brain connectivity (because of a difference between brain development and ontogenesis), but also wiring specificity between brain areas (AL and LH). Combining anatomical and physiological approaches, we demonstrate that scalable microcircuits within distinct glomeruli mediate the robust encoding of odor stimuli, while the different convergence of odor information onto LH neurons leads to differences in innate odor-driven behaviors in flies developed at different temperatures.

### Mechanisms for robust function in a synaptically enriched circuit

The analysis of neural connectivity of distinct glomeruli reveals two important consequences of developmental temperature. First, an overall increase in synaptic partners corresponds to a larger number and larger size of synapses (Brp enriched). This synaptic scaling is consistent with the larger calcium transients measured in some of the glomeruli in ORNs axons (Fig. 4). However, within all glomeruli a larger number of LNs is recruited at lower temperatures, likely driving higher inhibitory feedback. The balance between excitation and inhibition is key for the function of healthy brains and there is evidence that this balance is achieved developmentally during synapse formation^38^. In sensory systems, lateral inhibition sets the operational set point for stimulus encoding through a divisive normalization^39,40^. In the fly olfactory system, GABAergic inhibition is distributed at both pre- and post-synapses in ORN-uPN connections^41,42^. This agrees with our observation that in some glomeruli calcium transients are already compensated at the ORN axon terminals, while in others, odor response is only fully robust in second order neurons, after the first synaptic connection (Fig. 4). However, it remains unclear whether the scaling of inhibition is the sole mechanism that keeps responses temperature invariant. It is possible that the presynaptic scaling is also a homeostatic response to weaker post-synapses^43,44^. Overall, we conclude that the peripheral olfactory system is designed to compensate the temperature driven synaptic scaling and to keep odor information invariant to shifts in developmental temperature.

### Circuits for the modulation of odor preference downstream of the AL

Despite the robust odor representations within the AL, behavioral responses to appetitive odor cues are strongly affected by developmental temperature in two different assays (Fig. 3). Innate odor preference arises in the LH where uPNs target LHNs within a rather complex wiring logic^45,46^. Our analysis demonstrates that LHNs receive temperature dependent odor information from the antennal lobe through the differential wiring of uPNs (Fig. 5). It should be noted that the measured change in connectivity is only a lower bound, as only half of the uPNs were included in our analysis. Nonetheless, we provide evidence that the effect of developmental temperature is not restricted to a single LHN type. We further demonstrated that PD2b1 neurons, differentially wired to the AL, have different odor responses, and therefore LH output is not invariant across developmental temperatures. PD2a1/b1 neurons are required for memory retrieval^47^, but their role in innate behavior is unclear. They might be involved in approach towards appetitive cues, although their activation is not sufficient to drive behavior^46^. Importantly, PD2a1/b1 neurons might act synergistically or redundantly with other LHNs with temperature-dependent wiring, although the logic of these circuits remains poorly understood. We have further identified a change in wiring between ORNs and mPNs that target the LH and the protocerebrum. The function of this large population of neurons (>50% of the total number of PNs) is still unclear. Half of these neurons are predicted to be GABAergic^20^; GABAergic mPNs have modulatory effects on preference and discrimination^21,23,24^, and silencing inhibitory mPNs reduces approach towards appetitive odors^21^. The other half of mPNs is predicted to be cholinergic^20^ and their contribution to downstream odor processing or behavior is unknown. Further understanding of the LH circuit organization and mPN function is required to link the temperature-dependent plasticity to behavior. Nonetheless, we can conclude that the behavioral consequences of development at different temperature derive from a distributed effect on the organization of olfactory inputs into the LH, and not from a general enhanced sensitivity to odors within the peripheral sensory system.

### Metabolic constraints on the neural system development

As previously proposed by Kiral et al., an increased number of synapses at lower developmental temperature is the result of a difference between the scaling of circuit development and animal growth rates. Here we show that fly developmental times scale exponentially with temperature in agreement with proposed theories that assume a metabolic constraint to growth rate. We extended this theoretical framework to model the rate of neural growth and synapse formation. To do so, we made two assumptions. The first is that a different rate limiting reaction constraints neural development compared to ontogenesis (whole animal development). Such limiting rate could additionally vary across subcircuits or cell-types leading to differences in synaptic scaling across brain regions. In line with this, recent finding in zebrafish demonstrated that developmental temperature could have cell-type specific effects on the cell biology^48^. The second assumption we made is that neural/synaptic growth scales with the fractional power z=¾. This value was previously justified starting from geometrical considerations on the fractal nature of the distributions of resources throughout capillaries in a 3D volume^31^. Our data at fixed temperatures are consistent with any z<1, but the developmental outcome measured in presence of cycling temperatures is well predicted by a ¾ exponent^49^.The highly plausible assumptions, the success of the model in explaining our data and the compatibility with observations in another sensory circuit^16^, suggest that similar principles may apply to other circuits and organisms.

### Phenotypic variation and constrained pathways

Overall, our study suggests that phenotypic variation should be variably expected upon a temperature shift during development depending on the different metabolic constraints on growth imposed on different neural subcircuits. Synaptic scaling likely recruits partners based on their spatiotemporal availability. Clearly some connections in the brain are critical for survival and these have probably evolved to correctly wire no matter the temperature. Experiments done at extremely high temperatures reveal the backbone of the olfactory pathway, i.e. the wiring of ORNs onto a single uPN, a few mPNs and LNs (Fig. 2b). Studying these extreme conditions might reveal insights on evolutionary constrains on circuit design.

The brain has evolved many strategies to keep circuits’ function robust to environmental factors^50,51^. While such robustness holds true within some subcircuits (individual glomeruli in our study), it cannot be assumed to occur throughout the brain, in our case in the downstream pathways to the LH. Our study raises the question of whether variation in brain wiring is an evolutionary selected feature for adaptation to the environment, posing a new challenges for the understanding of brain development in poikilothermic animals.

## Acknowledgment

We thank Marion Silies for access to resources, Marion Silies, Filippo Calzolari and Sebastian Cachero for critical reading of the manuscript, Christopher Schnaitmann for help with *in vivo* imaging, Luisa F. Ramirez Ochoa for discussions of the model, Constantin Müller, Nancy Benja and Polina Krasnova for help with behavioral experiments, the biology workshop at JGU for developing the behavioral assay, Christoph Rickert for help with confocal imaging, Maria Ioannidou, Tristan Walter, Fritz Francisco and Angela Albi for help with video tracking, Sebastian Cachero and Greg Jefferis for sharing a new BAcTrace construct, David Deutsch for materials for the spherical treadmill, Robin Hiesinger, Carsten Duch, Bassem Hassan, members of the FOR5289 and members of the Silies lab for discussions. We further thank Sabine Schmitt, Simone Renner and Jonas Chojetzki for technical and administrative support.

## Author contribution

P.Z., L.B. and C.M. designed the study. P.Z. and L.B. performed and analyzed the anatomical experiments, L.B., S.C.B. and G.D.U. performed and analyzed *in vivo* calcium imaging, G.D.U., C.D. and S.C.B. set up, performed, and analyzed behavioral experiments, C.M. performed electrophysiology, analyzed data, developed the model, C.M. and L.B. wrote the manuscript and all authors edited it. This work was supported by the DFG grants MA7804/2-1 and MA7804/3-1 to C.M.

## Materials & Correspondence

Carlotta Martelli

## STAR Methods

### Key resources table

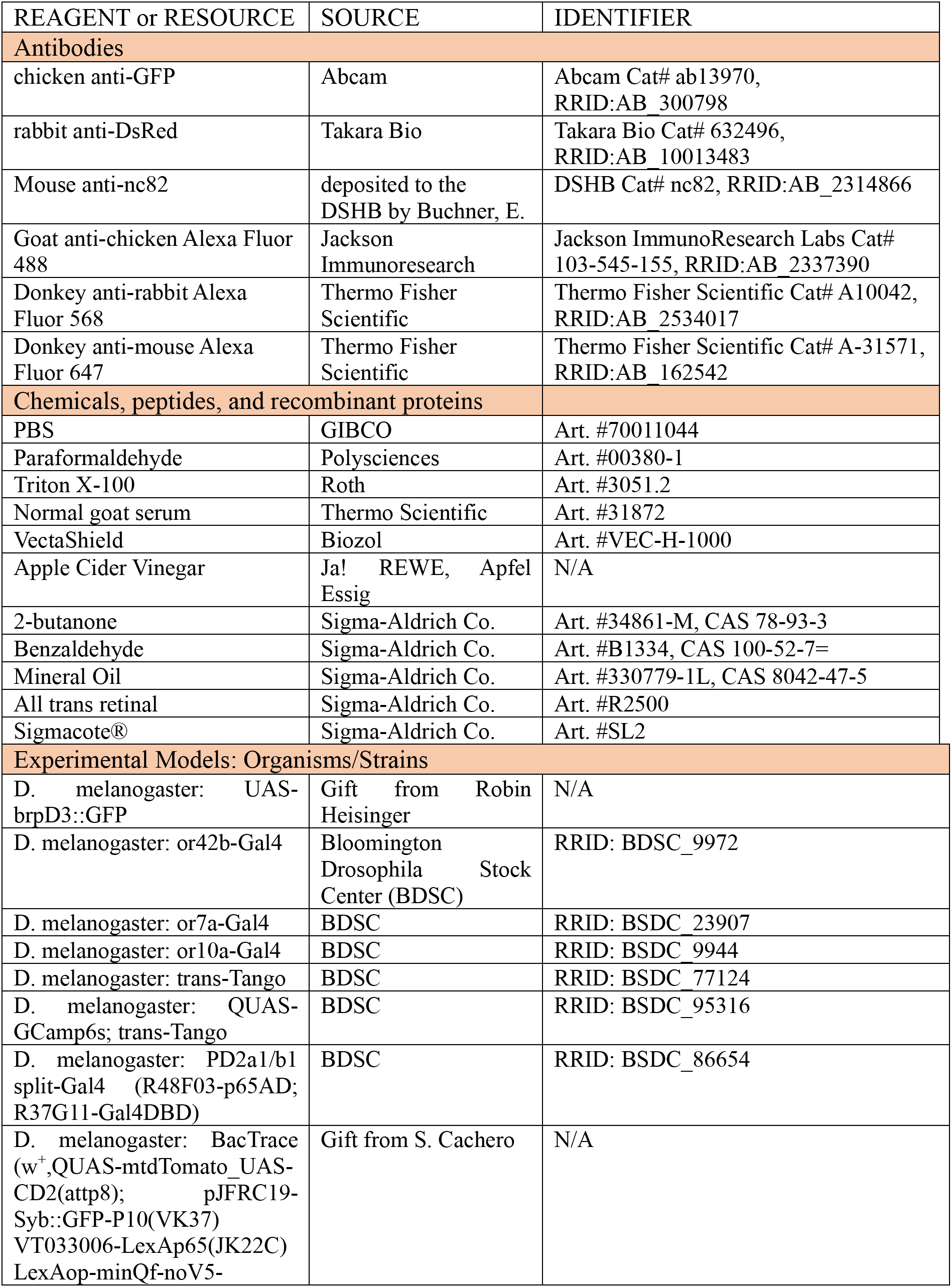
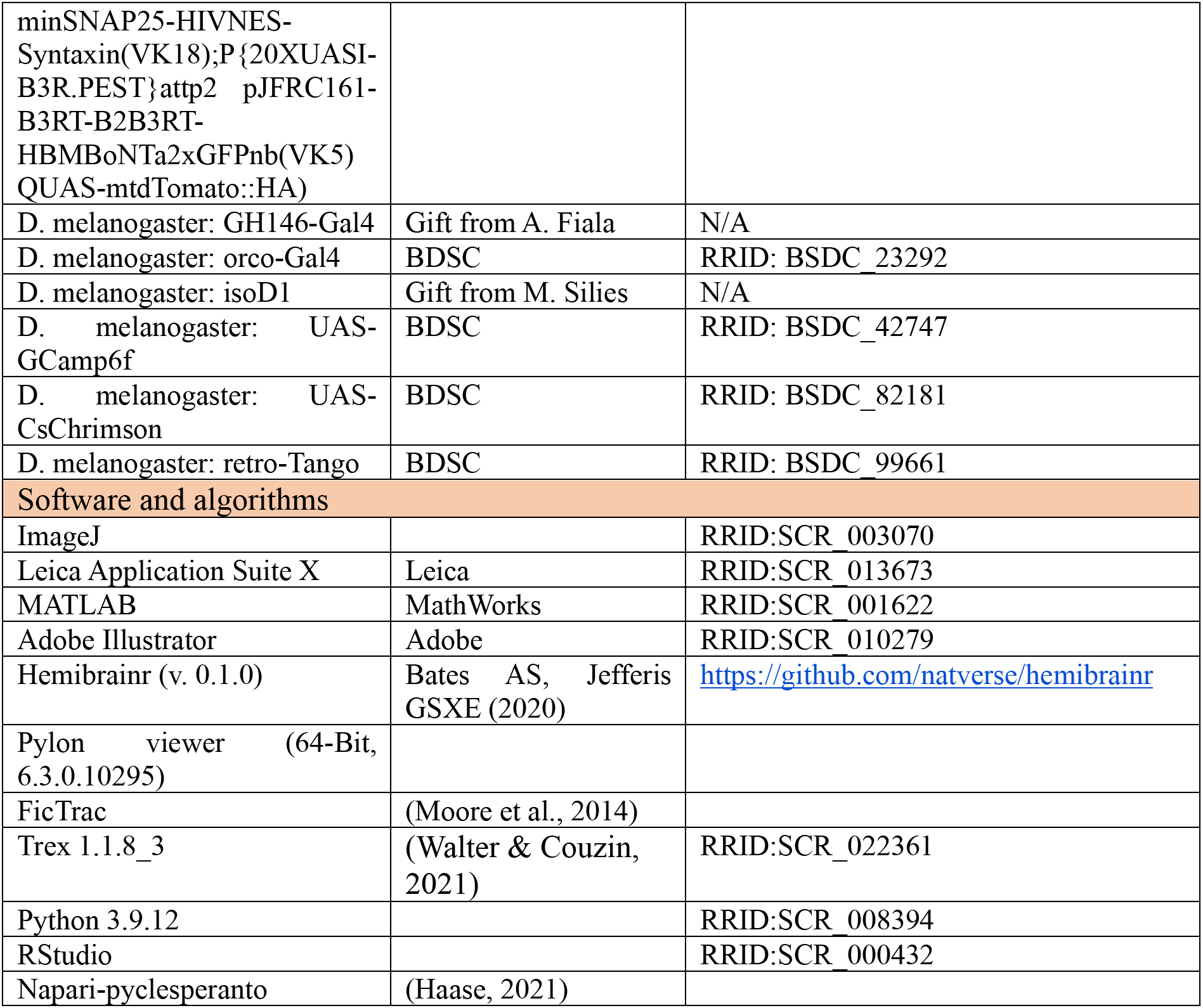

### Method Details

#### Experimental model / Fly husbandry

Flies were raised on standard molasses-based food, at 65% humidity and on controlled 12h:12h light-dark cycle. All flies were kept at 25°C as embryos and after eclosion. Flies were placed at different temperature (12°C, 18°C, 25°C and 31°C depending on the experiment) between the second instar larva stage (L2) and the end of metamorphosis. As the time of development depends on temperature, we call the end point of metamorphosis P-100%. For optogenetics experiments, flies were kept in standard molasses-based food with 1mM all-trans retinal (Sigma-Aldrich) in the dark for ≥ 72h before experiments. In all experiments only females were used. Exact genotypes are given in the table below.

| Name | Genotype | Figure |
| --- | --- | --- |
| or42b > trans-Tango | w+, UAS-myrGFP.QUAS-mtdTomato-3xHA; trans-Tango / CyO ; or42b-Gal4 / + | Fig. 1a-d, Fig. 1m, Fig. 2a-e |
| or42b > trans-Tango QUAS-Gcamp6s | w+ 10x QUAS-Gcamp6s; trans-Tango / UAS-CsChrimson; or42b-Gal4 / + | Fig. 1e-h Supp. Fig. 1a |
| or42b > brpShort GFP | ;+;UAS-BRP D3::GFP/or42b-Gal4 | Fig. 1i-j Supp. Fig. 1c |
| GH146 > GCamp6f | w+; UAS-Gcamp6f / GH146-Gal4 | Fig. 2n |
|  |  | Fig. 4d |
| Or42b > GFP | w+, UAS-myrGFP.QUAS-mtdTomato-3xHA;<br>trans-Tango / CyO ; or42b-Gal4 / + | Fig. 4b |
| orco > GCamp6f | w+; UAS-Gcamp6f / +; orco-Gal4 / + | Fig. 4a,c<br>Supp. Fig. 3 |
| PD2a1/b1 > BacTrace | w+, QUAS-mtdTomato,UAS-CD2/+; Syb::GFP,<br>VT033006-LexAp65, LexAop-minQf-noV5-<br>minSNAP25-HIVNES-Syntaxin/R48F03-<br>p65AD; UASI-B3R.PEST,B3RT-B2B3RT-<br>HBMBonTa2xGFPnb, QUAS-mtdTomato::HA /<br>R37G11-Gal4DBD | Fig. 5b-e<br>Supp. Fig. 4 |
| PD2a1 > Gcamp6f | w+; UAS-Gcamp6f / R48F03-p65AD ; R37G11-<br>Gal4DBD / + | Fig. 5j |
| AD1C1 > BacTrace | w+, QUAS-mtdTomato,UAS-CD2/+; Syb::GFP,<br>VT033006-LexAp65, LexAop-minQf-noV5-<br>minSNAP25-HIVNES-Syntaxin/R55C09-GAL4;<br>UASI-B3R.PEST,B3RT-B2B3RT-<br>HBMBonTa2xGFPnb, QUAS-mtdTomato::HA /<br>+ | Fig. 5f<br>Supp. Fig. 4 |
| AV4A1 > BacTrace | w+, QUAS-mtdTomato,UAS-CD2/+; Syb::GFP,<br>VT033006-LexAp65, LexAop-minQf-noV5-<br>minSNAP25-HIVNES-Syntaxin/ R34C08-<br>p65.AD; UASI-B3R.PEST,B3RT-B2B3RT-<br>HBMBonTa2xGFPnb, QUAS-mtdTomato::HA /<br>R22C12-GAL4DBD} | Fig. 5f<br>Supp. Fig. 4 |
| PD2a1 > retro-Tango | y w+ QUAS-mtdTomato-3xHA; retro-Tango / + ;<br>10xUAS-retro-Tango/ R48F03-p65AD; R37G11-<br>Gal4DBD / + | Fig. 5i |
| or7a > trans-Tango | w+, UAS-myrGFP.QUAS-mtdTomato-3xHA;<br>trans-Tango / or7a-Gal4 | Fig. 1k,n |
| or10a > trans-Tango | w+, UAS-myrGFP.QUAS-mtdTomato-3xHA;<br>trans-Tango / or10a-Gal4 | Fig. 1k,o |

#### Temperature cycle experiment

For temperature cycle experiments flies were kept at an 12h: 12h light-dark cycle, and either a temperature cycle/protocol of 20-28°C or 16-32°C, with 8h of temperature plateau and temperature changes of 2°C/h and 4°C/h, respectively.

#### Immunohistochemistry and confocal

Female flies (9-11 days post eclosion) were anesthetized with ice and then briefly submersed on ethanol 70%. Flies were dissected on cold Phosphate Buffer Saline (PBS) for no longer than 20 minutes and fixated for 50 minutes in 2% paraformaldehyde (PFA, Polysciences, diluted in PBS) rotating at room temperature. All subsequent incubation and washes were done while rotating, in the dark. Brains were washed three times in PBT (PBS with 0.5 % Triton x-100, Roth) for 15 min and then blocked for one hour in 5% normal goat serum (NGS, Thermo Scientific, in 0.3 % PBT). Samples were incubated in primary antibody mixture (chicken anti-GFP 1:1000; rabbit anti-DsRed 1:500; mouse anti-nc82 1:25) for 48hrs at 4°C, then washed three times in PBT and incubated in secondary antibody mix (goat anti-chicken Alexa Fluor 488; donkey anti-rabbit Alexa Fluor 568; donkey anti-mouse Alexa Fluor 647, all at 1:200) for 48hrs at 4°C. Finally, brains were washed three times in PBT and mounted in VectaShield (Biozol) antifading medium. Brains were imaged on a Leica SP8 microscope with a 20, 40 or 63x objective depending on the experiment.

After image acquisition, the number cell bodies of postsynaptic partners were manually counted using Fiji’s cell counter plugin. Cell body numbers were classified according to the position around the antennal lobe: dorsal, lateral or ventro-lateral (including both ventro-lateral and ventral clusters).

##### BrpShort analysis

Confocal images of individual DM1 glomeruli were processed using a custom code in python, with the package pyclesperanto (Haase, 2021). Images contained a brp-Short and nc82 channel. Both channels were pre-processed with Gaussian Blur (1.0, 1.0, 1.0) and top hat box (20.0, 20.0, 1.0). DM1-ORNs mask was made using the Brp-Short channel, by applying Voronoi Otsu Labeling and then merging the touching labels. For the whole DM1 glomerulus mask, the DM1-ORN volume was closed using the function closing labels. Single nc82 puncta were labeled by Voronoi Otsu Labeling and restricted to the DM1 glomerulus mask. Labels volume and fluorescence intensity were acquired using the statistics of labelled pixels from pyclesperanto.

#### *In vivo* calcium imaging

Flies developed at either 18°C or 25°C, 9-11 days post eclosion, were anesthetized on ice and mounted on a custom holder using UV-cured glue (Bondic). Saline solution (5mM Hepes, 130 mM NaCl, 5mM KCl, 2 mM MgCl_2_, 2mM CaCl_2_, 36 mM Saccharose – pH 7.3) was added. The cuticle covering the fly’s head, as well as obstructing trachea, were removed.

Functional imaging was done on an Investigator two-photon microscope (Bruker) coupled to a tunable laser (Spectraphysics Insight DS+) with a 25×/1.1 water-immersion objective (Nikon). Laser excitation was tuned to 920 nm, and less than 20 mW of excitation was delivered to the specimen. Emitted light passed through a SP680 short-pass filter, a 560 lpxr dichroic filter and a 525/70 filter. PMT gain was set to 850 V. The microscope was controlled with the PrairieView (5.4) software.

##### Optogenetics

For optogenetic activation light from a 625 nm diode was directed using an optic fiber to the fly’s antenna. The diode was controlled in flight-back mode from the imaging software, allowing simultaneous acquisition and excitation. The light stimulus protocol consisted of 5s series of light pulses, presented 5 times with intervals of 30s. Stimulus intensity in Fig. 1g was measured at the fly position with this protocol.

##### Odor delivery

Flies were exposed to a continuous clean air airflow (1L/min), in which either an odor stream (100mL/min) or a clean balancer airflow (100 mL/min) was redirected through a solenoid valve (LEE), so that the final airflow reaching the fly was around 1.1mL/min. For creating the gas dilutions four mass flow controllers were used (Analyt-MTC) and controlled using a custom MATLAB (MathWorks) script and an Arduino board. Odors were prepared as a liquid 5 mL 10^−2^ volumetric dilution in 20ml glass vials (2-butanone and benzaldehyde in mineral oil, apple cider vinegar in MiliQ Water). The final volumetric gas dilution used was 10^−5^. Odor stimulation consisted of three repetitions of a 5 s, with 30 s intervals in between.

### Electrophysiology

Single sensillum recordings were performed as previously described (Martelli et al., 2013) using a silver-chloride electrode and glass pipettes filled with sensillum lymph ringer. Electrical signals were amplified using an extracellular amplifier (EXT-02F-1, npi) with head stage (EXT-EH), bandpass filtered (300–5000 Hz), digitized at 20KHz using a NI board (NI-6212). Data were acquired with the matlab toolbox *kontroller* (Gorur-Shandilya et al., 2017) https://github.com/emonetlab/kontroller. Spikes were sorted using a custom MATLAB routine.

#### Odor delivery

Flies were exposed to a constant airflow (1L/min) and an odor stimulus was delivered by switching a 3-way solenoid valve that directed a secondary airflow (100 mL/min) through a Pasteur pipette as in (Martelli et al., 2013; Martelli & Fiala, 2019). The pipette contained a filter paper with 50ml odor dilution. Volumetric odor dilutions were prepared in either mineral oil or MiliQ Water. Stimuli were controlled by custom made software in MATLAB and arduino.

### Behavioral experiments

#### Spherical treadmill

Experiments were conducted at 32°C in a closed custom arena. The spherical treadmill consisted of a 15mm diameter polyurethane foam sphere (FR-7120 foam, General Plastics) floating on an air-column. The sphere was coated with two layers of classic wood glue (Ponal, 25% in water) and then a random non-uniform pattern was drawn using two layers of acrylic black paint (Black 3.0, Culture Hustle). All coats of paint were allowed to dry overnight. The odor delivery system was similar to the one described above for *in vivo* calcium imaging experiments, but with a differing airflow rate controlled by Alicat Scientific MFCs. Continuous clean airflow was 90 mL/min and both the odor and balancer airflows were 10 mL/min. Videos were acquired with a XIMEA xiQ video camera, placed 10cm from the treadmill. The treadmill ball was illuminated by a panel of 940nm LEDs (Solarox) and an extra LED on the air-column was visible in the video and turned on simultaneously to the odor stimulus to trigger the data acquisition.

For experiments, 9-10 days old female flies were cold anesthetized and secured to a needle at their thorax on the dorsal side using a UV-hardening glue (Bondic) and positioned on the sphere with the help of a 3D micromanipulator. Before starting to record, flies were acclimatized to walking on the sphere for 10-15 minutes with no stimulus being presented. Subsequently, video recording and the odor stimulation were started. Videos were acquired using the XIMEA CamTool software: the exposure was set to 10,725ms, the gain to 2.6Db and the framerate to 80fps. The odor stimulation was controlled through MATLAB by an ARDUINO UNO Rev3 and consisted of at least 19 repetitions of 5s long odor stimuli and 20s-long interval without odor.

Fly moving speed was calculated a posteriori from the video recordings using the open-source software library FicTrac (Moore et al., 2014). For this analysis, regions of interest (ROIs), ignored regions and the transformation from the camera’s frame of reference to the animal’s coordinate frame were defined with the help of a configuration utility provided in FicTrac. The transformation was only calculated once at the start of experiments, since the position of the camera was never altered, but the ROIs and the ignored regions were updated for each recording. After running FicTrac, an output file was generated containing the output data, which includes rotation and speed of the animal for each frame in the video. The video recordings were also analyzed in MATLAB in order to extract the timepoints where the air-column LED was on, which corresponded to the odor stimulus being presented to the fly.

Finally, the output data from Fictrac and the timepoints obtained from the video were analyzed in MATLAB so that the moving speed during and outside odor presentation could be quantified. Flies that had a basal walking speed lower than 2mm/s were discarded.

#### Free-walking assay

##### Arena design

A free walking area was contained in a thermally controlled black box (100×45cm×45cm) shielded from room light, fully closed with a frontal door, and equipped with a heating system and thermostat (H-TRONIC GmbH, Product ID: 1114430). The box is heated up by an air stream created by a fan (at the base of the box) and homogeneously distributed a diffuser. A blue LED stripe (470nm, Paulmann Licht GmbH, Product ID: 78979) was positioned around the walking arena to ensure stable illumination during experiments. Videos were recorded with a Basler Camera (Basler acA2040-90um) placed on the ceiling of the box and equipped with f12mm lens (Basler C10-1214-2M-S), using Pylon viewer (64-Bit, 6.3.0.10295).

The walking arena was 40.2 cm diameter and 2 cm height and composed of four stacked layers and three overlapping plates (glass or plexiglass). The bottom layer contained six holes, arranged at the corners of a hexagon, where 1.5mL glass vials (Fisherbrand 11565874) containing the test odor can be screwed in. On topo of this a Teflon® coated porous sheet (FIBERFLON GmbH & Co. KG, Product ID: 408.07 P) provides a walking surface for the flies hiding the door location. The mid arena layer consists in a sloped (at 11°, 5 cm length) ring that defines the accessible area. To sealing the walking arena, we used a glass plate coated with Sigmacote® (Sigma-Aldrich Co.) to prevent fly walking upside-down. This behavioral setup was built by the workshop of the Biology Department at Johannes Gutenberg Universität Mainz.

##### Experimental protocol

We tested female flies developed at 18°C or 25°C, 5-7 days post eclosion. One hour before the experiment, flies were transferred into a vial with only a small piece of filter paper soaked in water and kept at room temperature. For experiments carried at 32°C, the fly vials were incubated for 15min in a 32°C water bath.

To create an odor gradient inside the arena, five minutes before the start of each experiment, a 1.5ml glass vial containing 1mL of test odor was placed in one of the six possible odor positions in the behavioral setup. For each trial, a fresh odor vial was used, and the position was pseudo-randomized. Each trial consisted of 10-15 female flies exposed to either apple cider vinegar (10^−2^ in MilliQ water), 2-butanone (10^−2^ in mineral oil), or tested with empty vials.

Flies were gently pushed inside the arena using a custom fly transfer tube and the recording was immediately started. All experimental videos were recorded at 20 fps for 15 minutes and saved in mp4 format. At the end of each trial, the flies were removed and discarded. The initial condition was restored by removing the odor vial and the cover glass plate, replacing the Teflon® sheet with a clean one, and letting the whole system ventilate for 5 minutes.

##### Video processing and analysis

All required steps to pre-process the raw videos were done using the Python 3.9.12 distribution ANACONDA (Version 4.13.0). Scripts were written using Virtual Studio Code (Version 1.81.1). Recorded mp4 videos were processed and video tracked with the software TRex (Version 1.1.8_3) (Walter & Couzin, 2021). Fly trajectories were rotate to account for randomized odor position. The output files were analyzed using custom Python and MATLAB scripts. A threshold of 5cm was chosen to determine if the fly had located the odor source, variations of this threshold do not alter the results.

### EM analysis

We used the Hemibrain dataset (hemibrain:v.1.2.1) (Xu et al., 2020). For Fig.1l and Supp. Fig.1d, we considered all synapses from the or42b-ORNs of the left and right antenna within the DM1 glomerulus of the right hemisphere, which were selected by clustering the synapses bases on their 3d coordinates. The analysis was restricted to synapses of the right hemisphere, as postsynaptic partners are fully reconstructed only on this side. We calculated the number of synapses between each individual ORN and each individual post-synaptic neuron. Connections with less than 3 synapses from a single ORN were discarded. Moreover, postsynaptic neurons that received less than 10 total synapses were discarded. The same procedure was used for DL1 and DL5. The percentages of LNs and mPNs in Supp. Fig.1e are lower bounds, as calculated from the available annotations.

### Deriving synaptic scaling at different temperatures from a growth law with fractional power

To model ontogenetic growth, we follow the same approach of (Gillooly et al., 2002). We assume that growth scales as the 3/4 power of the mass *m* (see discussion in the main text and (West et al., 1999)) following the equation:

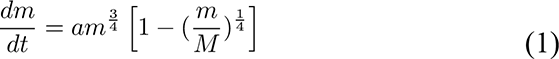

where *M* is the asymptotic mass and *a* is proportional to metabolic rate:

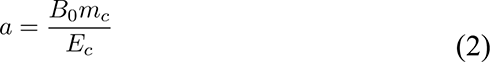

with *m_c_* the cell mass, *E_c_* the energy per cell and *B*_0_ is the normalization factor of the metabolic rate 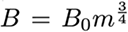 that scales proportionally to the Boltzmann factor: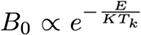 (*T_k_* is the temperature in Kelvin, *K* the Boltzmann constant and *E* the activation energy). Therefore 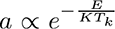. Following (Gillooly et al., 2002) who calculate *a* with respect to a reference temperature (the water freezing point *T_a_* = 273 K), and replacing *T* = *T_k_* – *T_a_*, we obtain:

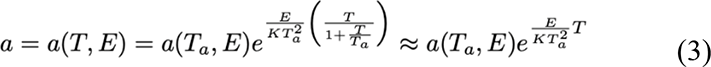

where now the temperature T is in °C. The last approximation takes in account the fact that the relevant temperatures do not exceed 32°C, therefore T/T_a_ is at most 0.1. This approximation leads to an error of about 10% on the exponential fits, but the quality of the model prediction remains unchanged. We keep the approximation for simplicity in the following calculations.

To find the relationship between developmental time and temperature, we integrate the growth equation (1) for *m*<<*M* (but see below):

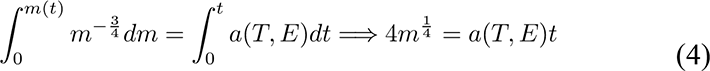

Equations (3) and (4) lead to the exponential relationship between developmental time t and temperature T proposed by (Gillooly et al., 2002):

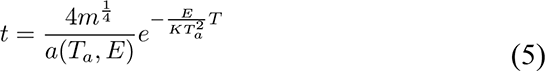

We calculate the fold change with respect to a reference temperature *T*_0_ = 25*°C* by assuming that development results in the same final mass:

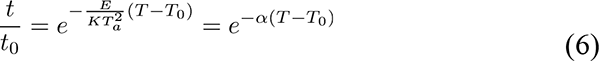

with 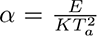. We use equation (5) to fit developmental times in Fig. 2h. This result remains the same if we use the general solution of equation (1) from (Gillooly et al., 2002)(relaxing the assumption *m*<<*M*) or if we integrate it from *m*(0) = *m_i_* instead of *m*(0) = 0.

We now assume that developmental time follows the exponential relationship in equation (6), while the wiring of the neural circuit is constrained by a different reaction rate with *E*^′^ *< E.* In modeling the growth of the neural system, we use *n* instead of the mass *m*, which can be intended as number of synapses, synaptic partners, or axonal branching. *n* follows a similar equation as (5) leading to:

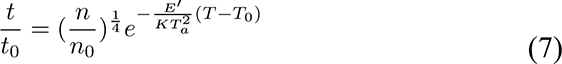

This results from an initial *n*(0) = 0, which is reasonable given the major pruning and re-growth of axons that happens during metamorphosis. Using equation (6) and (7):

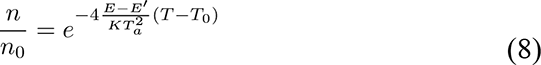

where we 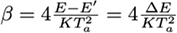. If Δ*E* = 0 then there should be no change in the number of synaptic partners. Also note that b can be larger than a 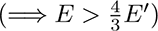 or smaller than a 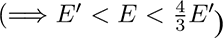. We use equation (8) to fit fold changes in number of synaptic partners in Fig. 2g.

To calculate developmental time of flies on temperature cycles with max and min temperatures *T*_1_ and *T*_2_. Here we simplify the cycling temperature protocol to step changes such that the final mass on temperature cycles results from development that occurs half of the time at *T*_1_ and half at *T*_2_:

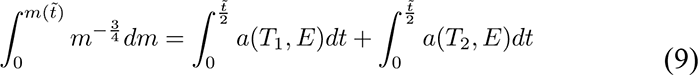

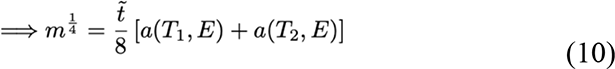

Here *t̃* indicates the developmental time on temperature cycles. Assuming an equal final mass, the fold change with respect to a fix temperature *T̅* is:

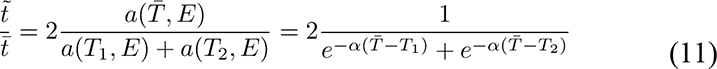

To calculate the fold change in synaptic connectivity, we use the same logic as before to derive:

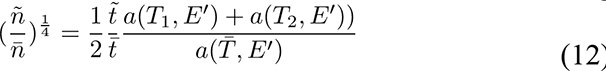

And using equation (11):

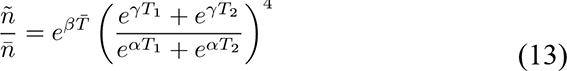

With 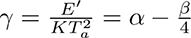. We use equation (13) to predict the number of synaptic partners in flies developed on periodic temperature cycles in Fig. 2k. The solutions (11) and (13) further simplify, if we take the mean temperature as the reference temperature 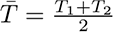 and 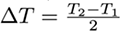 that is:

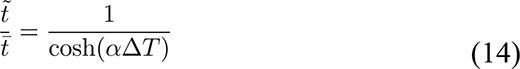

and

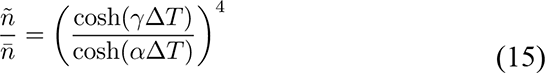

which show that the fold change in developmental time and connectivity scale inversely with the amplitude of the temperature cycles (as *γ* < *α*).

---

**Supplementary Figure 1.**
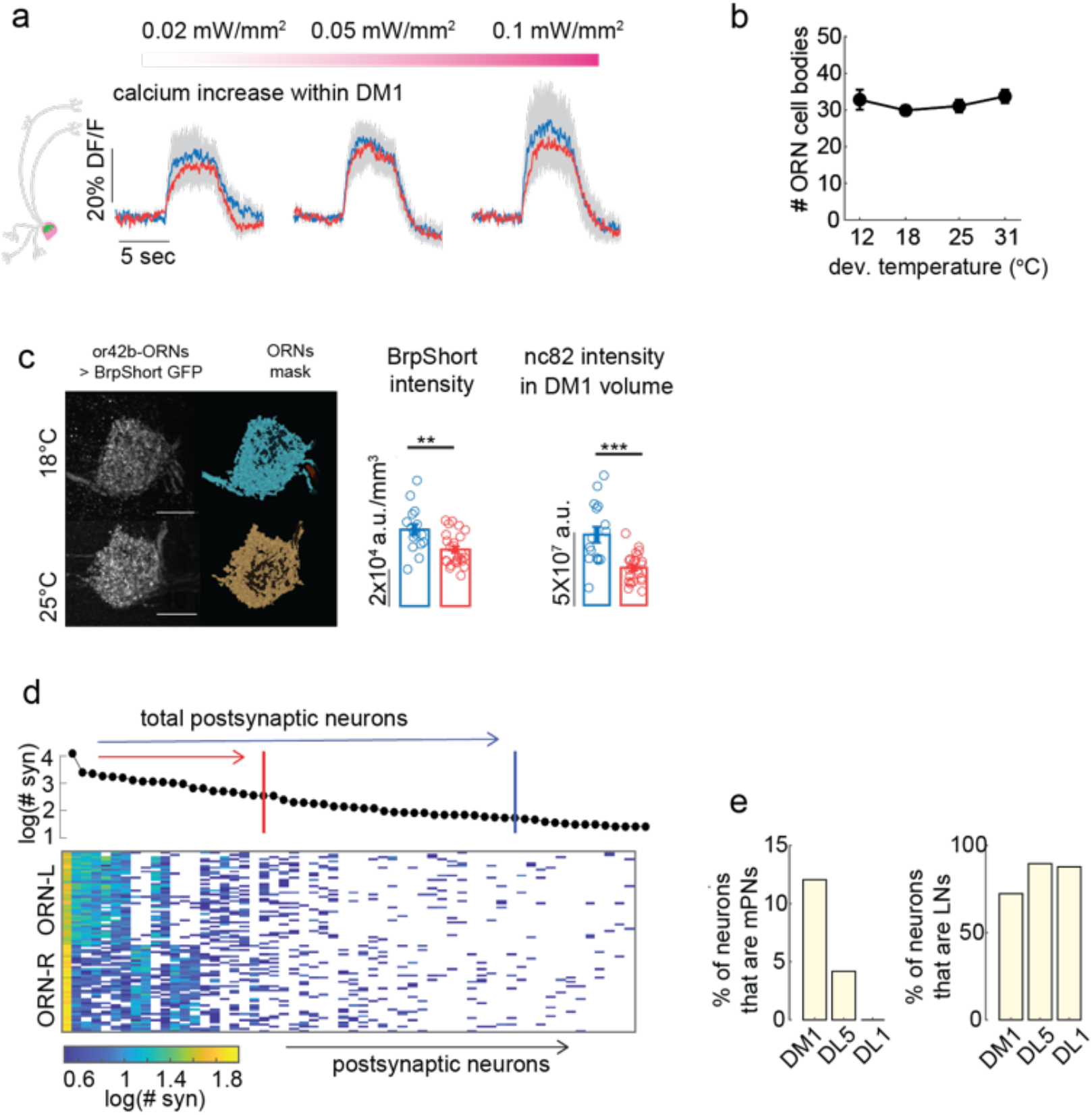
**a)** Optogenetic activation of or42-ORNs in flies developed at different temperatures. The usual trans-Tango reporter dsRed was replaced by GCaMP6s, and activity of presynaptic neurons induced by CsChrimson. Mean optogenetic response of DM1-postsynaptic partners was quantified in the glomerulus for three stimuli intensities. **b)** Mean number of ORNs cell bodies as a function of temperature (error bars indicate SEM). **c)** Left to right: Brp[Short]-GFP fluorescence expressed in or42b-ORNs and glomerulus 3D mask reconstructed from it for the two temperatures. Quantification of Brp^[Short]^-GFP and nc82 puncta intensity within the whole glomerulus volume (error bars indicate SEM, n = 16 hemibrains at 18°C and n = 25 at 25°C, Kruskal-Wallis test, **p<10^−2^, ***p<10^−3^). **d)** Bottom: connectivity matrix from hemibrain EM data illustrating the number of synapses made by DM1-ORNs from both antennae on postsynaptic partners in the right hemisphere. Post-synaptic partners on the x-axis are ranked by the total number of input synapses from these ORNs, as quantified in the top plot. Color scale indicates the number of synapses in log scale. The red and blue bar indicate the mean number of postsynaptic partners counted within one hemisphere in the trans-Tango experiment. **e)** Percentage of postsynaptic neurons that have been annotated in the Hemibrain dataset as multiglomerular projection neurons (mPNs, left) and local neurons (LNs, right).

**Supplementary Figure 2.**
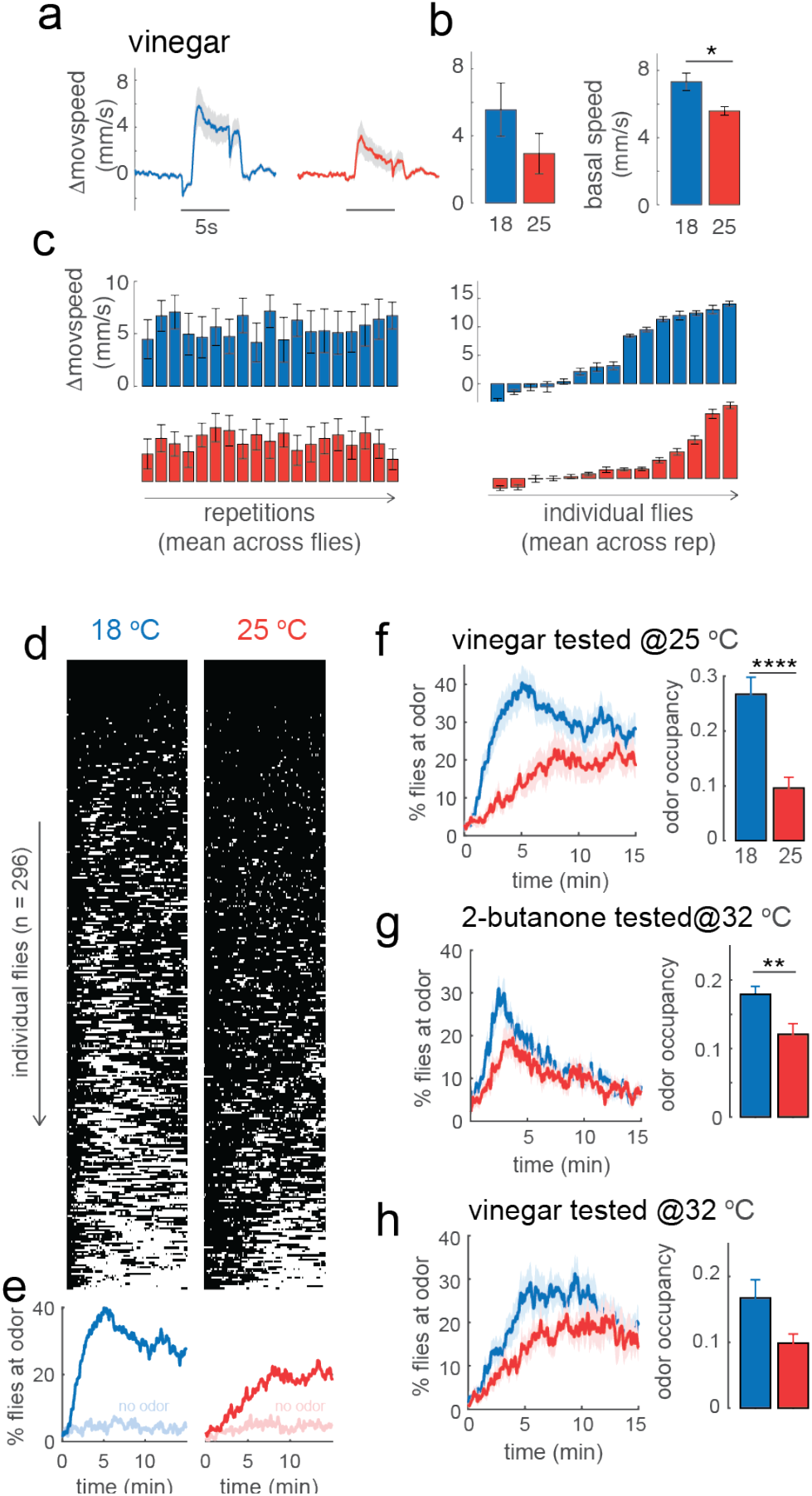
**a-c)** Response to vinegar on the spherical treadmill setup for flies developed at 18°C (blue) and 25°C (red). **a)** Odor response calculated as change in moving speed as a function of time in response to 5s stimulation; bar plot: mean change in moving speed within the first 2s of stimulation (p =0.1, n=14,13). **b)** Basal walking speed is estimated within 3s before stimulus onset (p = 0.02). **c)** Mean response to each consecutive stimuli repetitions averaged across all flies (right), and mean response of each individual fly across repetitions, showing large variation across individuals (left). **d-h)** Behavioral response in the free walking assay. **d)** Binary maps showing individual flies located at the odor source (<5 cm, white) as a function of time and **e)** percentage of flies at the odor (vinegar). An empty odor vial was used as control in an independent set of experiments (shaded curves). **f)** Average percentage of flies at the odor source and odor occupancy (bar plot), for vinegar tested at 25°C (p<10^−4^, n=21,20). **g)** same as in **f)** for a testing temperature of 32°C for 2-butanone (p=0.006, n=20,20) and **e)** vinegar (p=0.09, n=19,18). Shaded area and error bars indicate SEM.

**Supplementary Figure 3.**
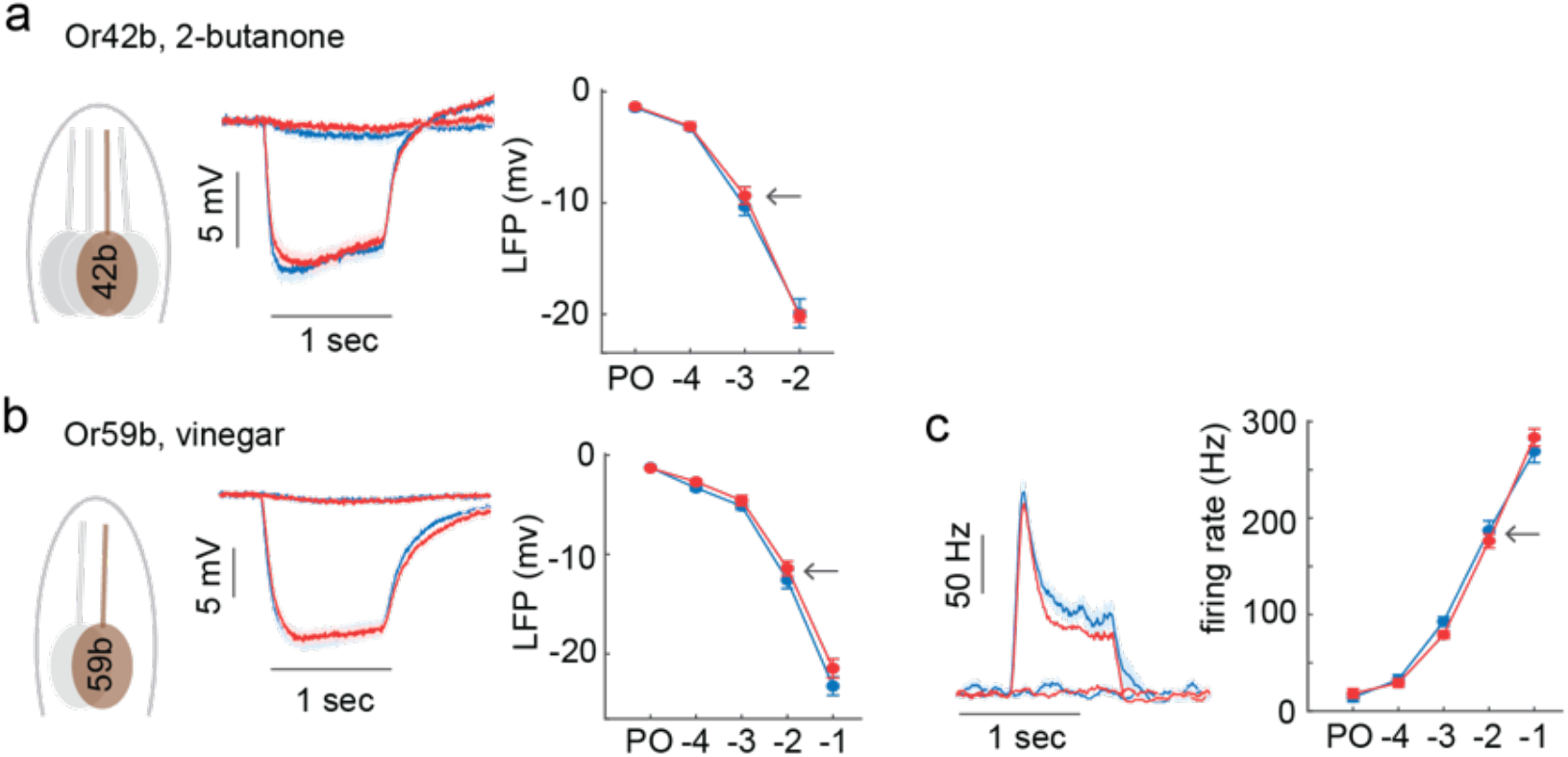
**a)** Left to right: Single sensillum recordings from the ab1 sensillum containing a single or42b-ORN. Mean response of the LFP to a 1s pulse of 2-butanone at 10^−3^ dilution and Paraffin Oil (PO) control. Amplitude of the LFP calculated as the maximum drop for each odor concentration. **b)** Same as a for the ab2 sensillum containing one or59b-ORN. **c)** Firing rate response of ab2 to a 1s pulse of 2-butanone at 10^−3^ dilution and PO control calculated in 100ms sliding window. Right: mean peak response for each concentration tested.

**Supplementary Figure 4.**
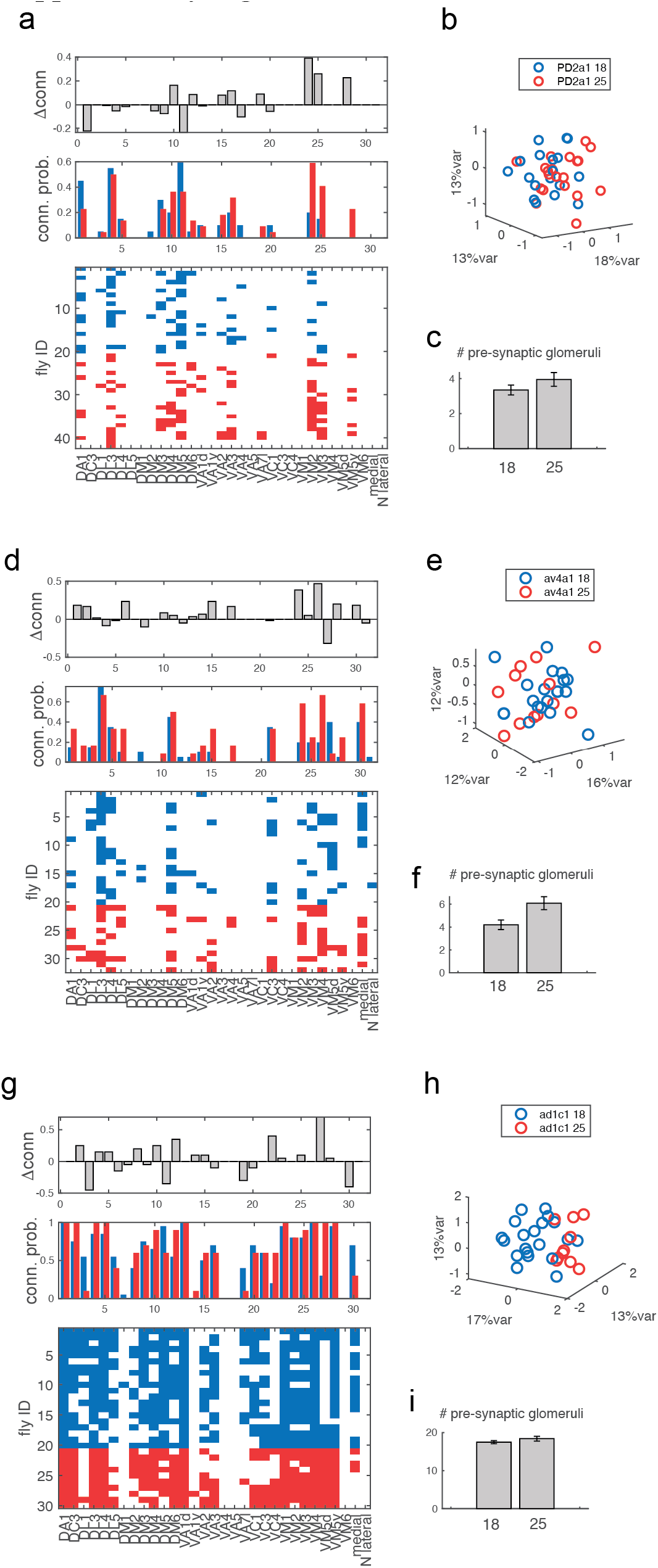
Connectivity analysis for three LH neurons. **a)** Bottom: binary connectivity matrix for pb2a1/b1 as in <u>Fig. 5c-d-e of the main text</u>. Middle: Connection probability to each glomerulus. Top: difference in connectivity between 25°C and 18°C. PCA of connectivity matrix: each dot indicates a single hemibrain**. c)** Mean number of pre-synaptic glomeruli for the LHN in each hemibrain. **d-e-f)** and **g-h-i)** same for av4a1 and ad1c1 referring to Fig. 5f <u>in the main text</u>.

